# Mitigation of Aβ_25-35_-Induced Cognitive Deficits in C57BL/6 Mice via Thermal Cycling Stimulation Employing Focused Ultrasound

**DOI:** 10.64898/2025.12.10.693347

**Authors:** Guan-Bo Lin, Hsu-Hsiang Liu, Yu-Yi Kuo, You-Ming Chen, Fang-Tzu Hsu, Yu-Wei Wang, Yi Kung, Chien Ching, Chih-Yu Chao

**Affiliations:** Department of Physics, Laboratory for Medical Physics & Biomedical Engineering, National Taiwan University, Taipei 106319, Taiwan; Molecular Imaging Center, National Taiwan University College of Medicine, Taipei 100233, Taiwan; Graduate Institute of Applied Physics, Biophysics Division, National Taiwan University, Taipei 106319, Taiwan

**Author notes:** Correspondence: Chih-Yu Chao, Department of Physics, Lab for Medical Physics and Biomedical Engineering, National Taiwan University, No 1, Sec 4, Roosevelt Rd., Taipei 106319, Taiwan, Republic of China, (CYC). (G.B.L.); (H.H.L.); (Y.Y.K.); (Y.M.C.); (F.T.H.); (Y.W.W.); (Y.K.); (C.C.); (C.Y.C.).

## Abstract

Hyperthermia (HT) is recognized across various medical disciplines for its capacity to modulate specific protein expressions. In efforts to improve Alzheimer’s disease (AD), HT has the potential to regulate heat shock proteins (HSPs) and antioxidant enzymes, which helps decrease the aberrant accumulation of β-amyloid (Aβ) protein and oxidative stress. Nonetheless, the precise delivery of mild hyperthermia to the brain remains a significant challenge. To apply mild hyperthermia targeted to the brain and evaluate its impact on cognitive improvement, this study used focused ultrasound (FUS) to administer localized mild hyperthermia to the brains of AD mouse induced by intracerebroventricular (i.c.v.) injection of Aβ_25-35_. For considerations of safety and therapeutic efficacy, a thermal cycling-hyperthermia (TC-HT) protocol was adapted into a focused ultrasound-mediated thermal cycling stimulation (FUS-TCS), which was compared with the continuous focused ultrasound-mediated hyperthermia stimulation (FUS-HTS). The findings revealed that the FUS-TCS treatment group exhibited a significant improvement in cognitive performance, as evidenced by enhanced outcomes in the Y-maze and novel object recognition (NOR) tests. Furthermore, this group demonstrated increased expression of Aβ-degrading enzymes and antioxidant proteins, including heat shock protein 70 (HSP70), neprilysin (NEP), insulin degrading enzyme (IDE), sirtuin 1 (SIRT1), and superoxide dismutase 2 (SOD2). These results suggest that localized mild hyperthermia targeting the brain using FUS-TCS treatment represents a promising strategy for ameliorating cognitive deficits associated with AD.

## Introduction

Hyperthermia (HT) represents a non-pharmacological approach to modulating biological responses by inducing alterations in cellular signaling pathways through temperature variations [1–3]. Various modalities of hyperthermia have been extensively applied in the treatment of diverse conditions, including muscle pain, arthritis, inflammatory disorders, and cancer [3–5]. In the realm of neurodegenerative diseases, hyperthermia has emerged as a potential therapeutic strategy for health improvement [6,7]. Alzheimer’s disease (AD), the most prevalent neurodegenerative disorder, is characterized by progressive cognitive decline and is experiencing a rising incidence globally. The aberrant accumulation of β-amyloid (Aβ) peptides in the brain is widely recognized as a major pathogenic factor in AD, contributing to oxidative stress, neuronal injury, and subsequent cognitive deficits [8–10]. Consequently, interventions targeting the mitigation of oxidative stress and the reduction of pathological Aβ depositions have garnered significant research interest [10–14].

Several protective proteins, which can be upregulated in response to hyperthermia, may underlie its capacity to mitigate Aβ accumulation and oxidative stress. Among these protective proteins, heat shock proteins (HSPs), particularly heat shock protein 70 (HSP70), are well-characterized molecular chaperones that are inducible by thermal stimulation. HSP70 contributes to the maintenance of protein homeostasis by preventing Aβ misfolding and aggregation, thereby facilitating its clearance [15,16]. Additionally, certain proteins known as Aβ-degrading enzymes, including neprilysin (NEP) and insulin-degrading enzyme (IDE), are implicated in the degradation and removal of Aβ, with their upregulation potentially contributing to the amelioration of AD pathology [12,17]. In addition to targeting Aβ, the regulation of oxidative stress represents another critical focus in AD pathogenesis. Proteins such as sirtuin 1 (SIRT1) and superoxide dismutase 2 (SOD2) play pivotal roles in modulating oxidative stress levels, thereby influencing the progression of AD [18,19]. Notably, several of the aforementioned protective proteins, including HSP70 and IDE, can be upregulated by hyperthermia, suggesting mechanistic pathways through which hyperthermia may confer neuroprotection and cognitive benefits [20–22]. These findings offer valuable insights into potential strategies for the prevention or mitigation of AD. Moreover, a Finnish study suggests that frequent use of Finnish saunas, particularly at moderate to high frequencies, may be associated with a reduced risk of developing AD [23]. Collectively, these observations position hyperthermia as a promising intervention capable of modulating key protein pathways implicated in AD, thereby offering potential therapeutic benefits.

Although these findings are encouraging, the majority of previous investigations have concentrated on in vitro experiments or whole-body hyperthermia techniques. The advancement of methodologies for the precise and safe application of hyperthermia to the brain remains underdeveloped, posing a significant obstacle to clinical translation. Focused ultrasound (FUS) offers a potential solution to this problem by directing ultrasound waves to a confined focal region, thereby minimizing effects on adjacent tissues. Clinically, high-intensity focused ultrasound (HIFU) enables noninvasive ablation of solid tumors by elevating focal temperatures above 55 °C, sometimes reaching 65 °C or even 80 °C, resulting in coagulative necrosis [24–26]. Moreover, HIFU is employed in medical aesthetics to target the deep dermis and superficial musculoaponeurotic system, where focal heating to approximately 60–70 °C induces collagen denaturation and contraction, facilitating skin tightening and rejuvenation [27,28]. In the context of brain applications, prior research has primarily utilized low-energy FUS for neural modulation or employed microbubbles to transiently disrupt the blood-brain barrier (BBB) to enhance drug delivery, rather than for thermal effects [29,30]. Different from both high-temperature HIFU and non-thermal low-energy FUS, we propose the use of FUS to induce mild hyperthermia in the brain through moderately elevated temperatures. Compared to whole-body hyperthermia, localized brain warming, as demonstrated in studies targeting brain tumors to improve drug delivery, allows for more precise stimulation of specific brain regions while limiting thermal exposure to surrounding tissues [31,32]. Nonetheless, prolonged exposure to elevated temperatures can exert thermotoxic effects on neural cells [33,34], and investigations into the neuroprotective potential of mild brain hyperthermia for neuroprotection remain scarce. Therefore, establishing a tolerable, effective, and safe approach for targeted brain hyperthermia is of critical importance for advancing AD research.

In our previous investigations, we introduced a novel therapeutic approach termed thermal cycling-hyperthermia (TC-HT), which demonstrated neuroprotective effects in vitro and was shown to be more efficacious than conventional hyperthermia (HT) [35,36]. Furthermore, in an animal study, we administered heat energy to the heads of mice using polyimide thermofoil and observed that intermittent heat treatment via TC-HT significantly improved cognitive function compared to continuous HT [37]. Building upon this prior external head-heating technique, the present study employed focused ultrasound-mediated thermal cycling stimulation (FUS-TCS) to deliver thermal energy directly to specific brain regions. The safety profile of FUS-TCS was assessed in comparison to continuous focused ultrasound-mediated hyperthermia stimulation (FUS-HTS). Cognitive performance in C57BL/6 mice exhibiting Aβ_25–35_–induced impairments was evaluated following FUS-TCS treatment using the Y-maze and novel object recognition (NOR) tests. Concurrently, we examined the expression of heat- and stress-responsive proteins implicated in Aβ clearance, including HSP70, IDE, and NEP, as well as antioxidant-related proteins such as SIRT1 and SOD2, which play critical roles in oxidative stress regulation. The results indicate that FUS-TCS may mitigate Aβ-induced cognitive deficits and modulate key proteins involved in AD pathology, thereby providing a promising framework for further preclinical evaluation and therapeutic development.

## Results

### Temperature assessment on the phantom during FUS-TCS application

To demonstrate the thermal distribution generated by FUS, a thermochromic phantom was employed to visualize temperature-dependent responses. The phantom comprised a transparent section and a white region containing thermochromic powder, facilitating the visual detection of localized heating through a color change occurring near 40.0 °C. As shown in Figure 1A, when the FUS transducer operated at 1.2 MHz, the thermochromic area appeared approximately 8 mm away from the transducer aperture, with a diameter of roughly 2–3 mm. On the other hand, at 1.8 MHz, the color change was observed approximately 15 mm from the aperture, with the thermochromic area measuring about 1 mm in diameter, as shown in Figure 1B. These findings indicate that FUS can deliver thermal energy precisely to a targeted area, and that varying the operating frequency produces different focal zones of energy deposition. To ensure mild hyperthermia within the targeted brain region of mice in subsequent experiments, an operating frequency of 1.2 MHz was chosen to better encompass the mouse brain. Before applying FUS-TCS to the mice, we measured the temperature profile in the phantom as shown in Figure 1C. During the 3-min high-temperature phase of FUS-TCS, the heating region’s temperature was maintained at 40.0 ± 0.5 °C, followed by a rapid drop to about 34.0 °C during the subsequent 1-min resting phase. Overall, the phantom results validated the selection of FUS-TCS parameters that provide consistent and reliable thermal energy delivery for the subsequent animal experiments.

**Figure 1.**
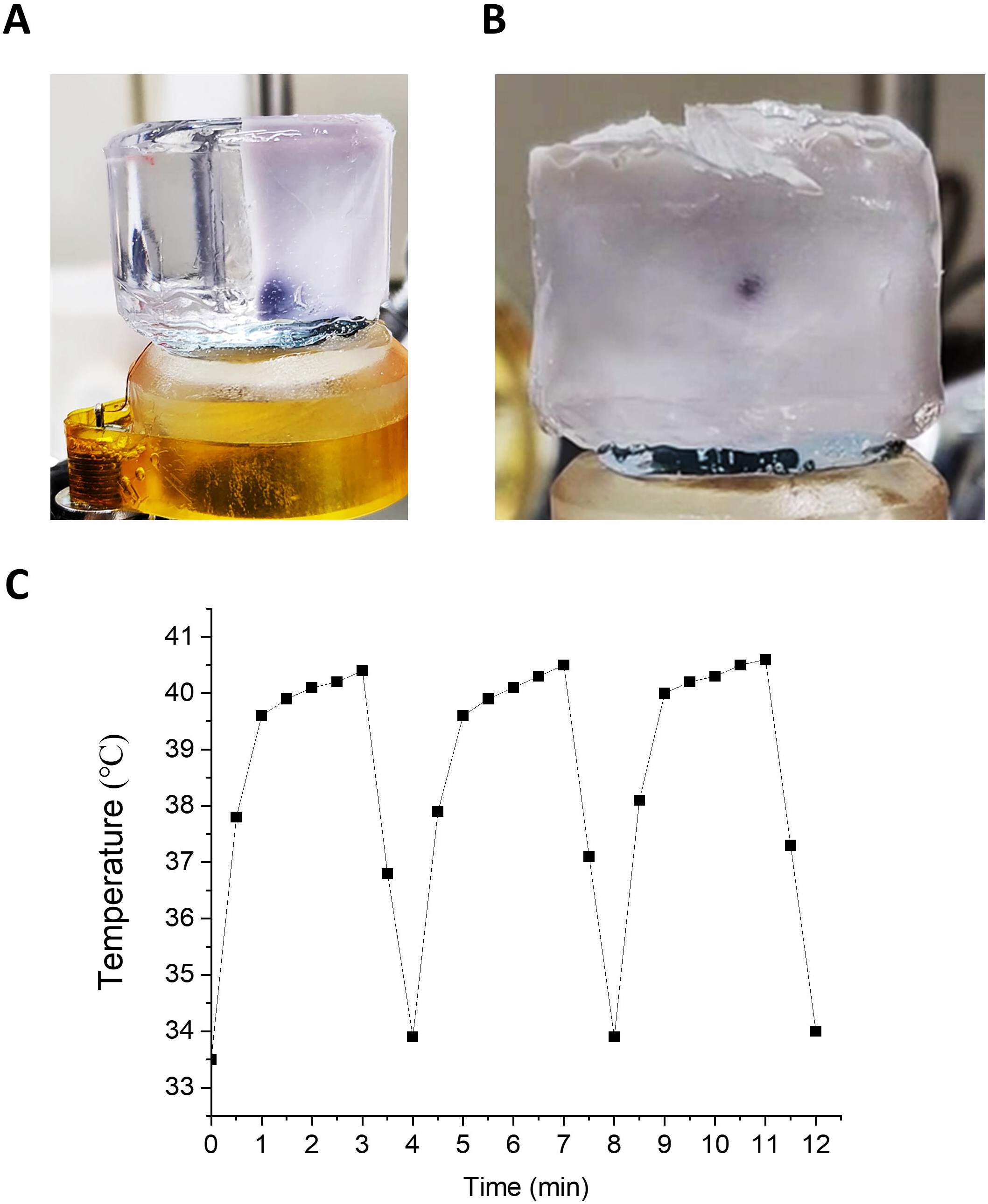
Representative thermochromic responses and temperature measurements in the phantom during FUS exposure. Colorimetric changes were observed in the thermochromic phantom following FUS application at frequencies of (A) 1.2 MHz and (B) 1.8 MHz. (C) The temperature profile recorded in the phantom during FUS-TCS application at 1.2 MHz is presented. The white region of the phantom was made from an acrylamide-based hydrogel formulation containing thermochromic powder, which exhibits a distinct visible color change at 40.0 °C. Temperature measurements were obtained via a K-type thermocouple placed at the geometric center of the thermal focal zone.

### In vivo mice brain temperature monitoring and experimental schedule

Before performing further experiments, we assessed the temperature profiles in the mouse brain during FUS exposure to verify the delivery of controlled mild hyperthermia. Temperature measurements were taken under anesthesia using a K-type thermocouple inserted directly into the mouse brain. As shown in Figure 2A, the FUS-TCS protocol involved a 10-cycle repeated heating process. Each cycle comprised a high-temperature interval lasting 3 min, during which the temperature was maintained at 39 ± 1 °C with peak temperatures reaching approximately 40 °C, followed by a 1-min passive cooling interval where the temperature dropped rapidly. For the FUS-HTS application, the temperature was also held at 39 ± 1 °C for 30 min. These findings indicate that mild hyperthermia can be precisely regulated through FUS application. After confirming the thermal stability, subsequent animal experiments were performed according to a predetermined schedule. As illustrated in Figure 2B, Aβ solution was administered via i.c.v. injection into the brains of mice in the Aβ, FUS-TCS, and FUS-HTS groups on day 0. The respective thermal treatments were applied to the FUS-TCS and FUS-HTS groups on days 4, 8, and 12. Behavioral assessments were subsequently conducted, including spontaneous alternation in the Y-maze on day 13 to evaluate locomotor activity and short-term spatial working memory. Additionally, NOR tests were performed on days 14 and 15 to assess short-term (1 h) and long-term (24 h) memory, respectively. These behavioral tests take advantage of mice’s natural tendency to explore new environments or unfamiliar objects [38–41].

**Figure 2.**
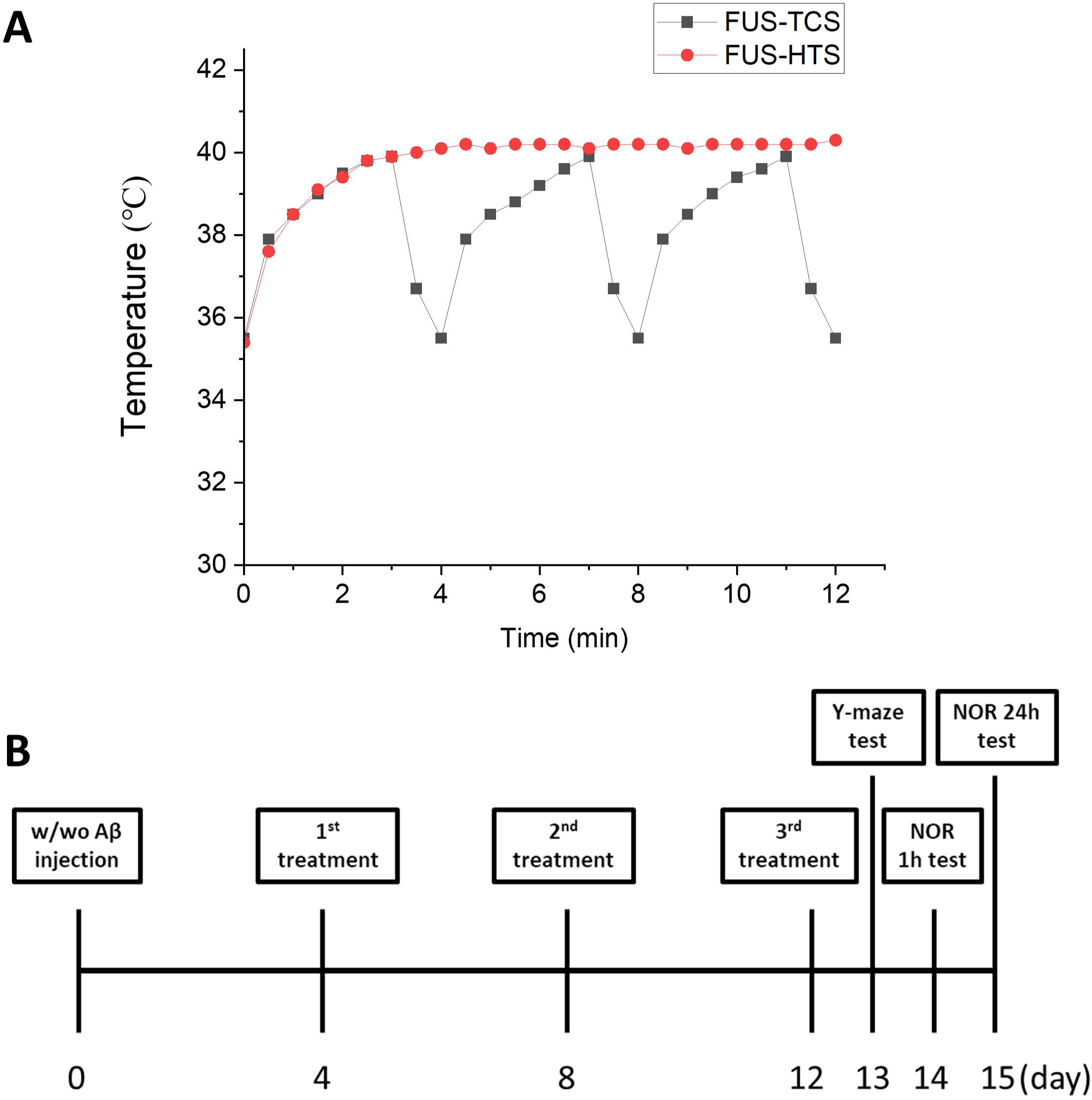
Temperature monitoring in the mouse brain during FUS stimulation and the experimental timeline. (A) Representative temperature recordings obtained from the mouse brain during FUS-TCS and continuous FUS-HTS treatments. Measurements were taken under anesthesia using a K-type thermocouple inserted through a small cranial opening. The temperature profiles illustrate the stable temperatures reached during FUS application, with FUS-HTS held around 40 °C and FUS-TCS varying between 35 and 40 °C. The brain temperature reached the target range within the initial TC-HT cycle and was sustained throughout subsequent cycles. (B) Diagram outlining the experimental timeline. Mice were sacrificed for brain tissue collection 24 h after the NOR test on day 15.

### Effects of thermal treatments on behavioral performance in mice

Before performing cognitive tests on AD mice, the safety profiles of two hyperthermic stimulation modalities, FUS-TCS and FUS-HTS, were evaluated. As shown in Figure 3A, there were no significant differences in the number of arm entries during the Y-maze test between the two thermal stimulation groups, suggesting that these treatments did not influence the mice’s locomotor activity. However, the FUS-HTS group demonstrated a significantly reduced spontaneous alternation index (51.0%) relative to the control (60.4%), whereas the FUS-TCS group did not differ significantly from the control group (Figure 3B). Furthermore, Figure 3C and D show that the discrimination index in the NOR test was reduced in the FUS-HTS group at both short-term (1 h) and long-term (24 h) intertrial intervals, decreasing from 63.1% to 42.1% at 1 h test and from 61.7% to 42.6% at 24 h test, respectively, compared to controls. In contrast, the FUS-TCS group maintained discrimination indices similar to the control group at both time points, indicating preserved cognitive function. These findings suggest that continuous thermal stimulation via FUS-HTS may impair cognitive performance, whereas intermittent mild hyperthermia induced by FUS-TCS appears to preserve cognitive function. Therefore, FUS-TCS was chosen for subsequent experimental investigations in AD mice.

**Figure 3.**
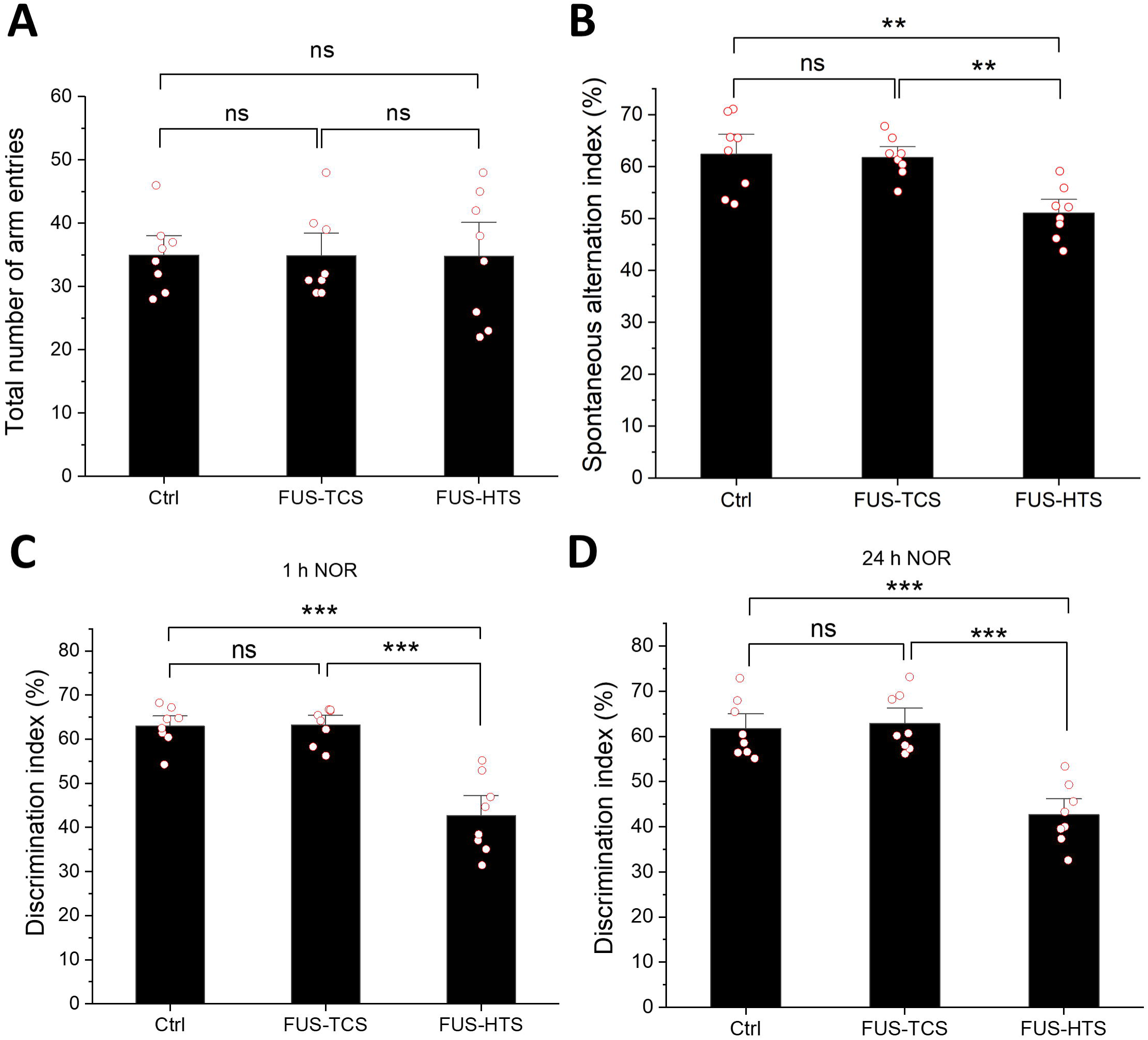
Effects of FUS-TCS and FUS-HTS treatments on behavioral performance in mice. Behavioral performance following different thermal treatments was assessed by (A) total number of arm entries and (B) spontaneous alternation index in the Y-maze test, as well as (C) discrimination index at 1 h and (D) 24 h intertrial intervals in the NOR test. Data are expressed as mean ± standard error of the mean (SEM). Statistical analysis was performed using one-way ANOVA followed by Tukey’s post-hoc test. (n = 8 in each group). Significant differences between groups are denoted as *P < 0.05, **P < 0.01, while non-significant differences are indicated as ns.

### FUS-TCS treatment alleviates Aβ-induced cognitive deficits in mice

To investigate the efficacy of mild hyperthermia induced by FUS-TCS in modulating cognitive deficits triggered by Aβ, behavioral assessments including the Y-maze and NOR tests were conducted. As shown in Figure 4A, there was no significant difference in the total number of arm entries among the control, Aβ, and Aβ+FUS-TCS groups, indicating similar levels of locomotor activity. However, Figure 4B demonstrates that mice administered with Aβ had a spontaneous alternation index that dropped from an average of 63.0% in the control group to 50.4%, indicating impaired working memory. Importantly, FUS-TCS treatment improved the spontaneous alternation index reduced by Aβ, bringing it close to the control level. Furthermore, Figure 4C demonstrates that Aβ-injected mice had a lower discrimination index at the 1 h intertrial interval in the NOR test, decreasing from 60.8% to 28.2%. In contrast, the Aβ+FUS-TCS group displayed an improved discrimination index of 55.8%, which was not significantly different from control. Similarly, at the 24 h intertrial interval in the NOR test (Figure 4D), the Aβ group exhibited a pronounced reduction in discrimination index (12.4%), whereas the application of FUS-TCS in Aβ-treated mice resulted in a substantial recovery to 56.2%, comparable to the control group. Overall, these findings suggest that FUS-TCS effectively mitigates Aβ-induced recognition memory deficits. Building upon these behavioral outcomes, we further explored whether FUS-TCS affects molecular pathways involved in Aβ clearance and antioxidant defense mechanisms.

**Figure 4.**
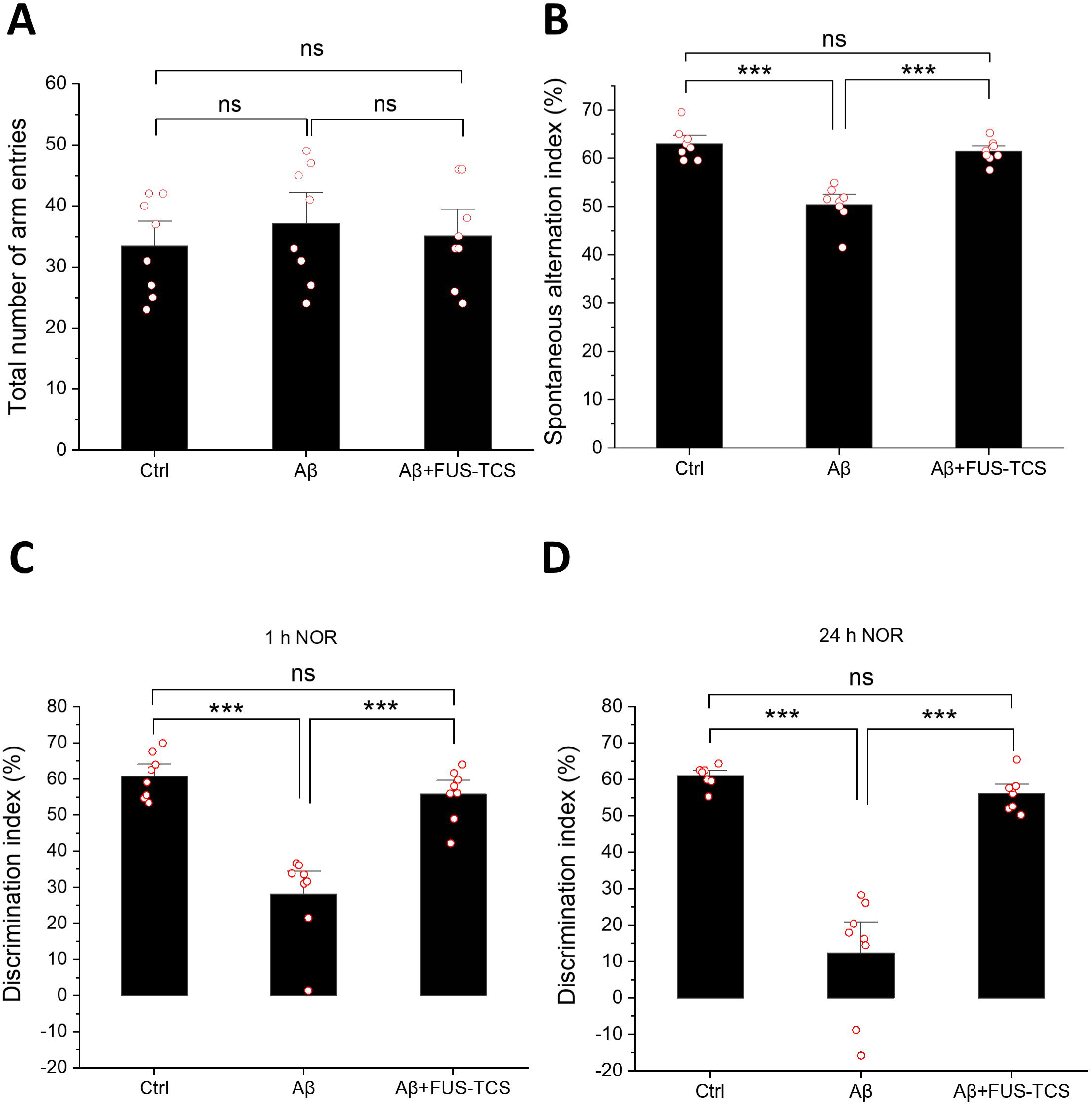
FUS-TCS mitigates Aβ-induced cognitive deficits in mice. Behavioral tests were conducted to assess locomotor activity and cognitive function among control, Aβ, and Aβ + FUS-TCS groups. The Y-maze test measured (A) total arm entries and (B) spontaneous alternation index, while the NOR test evaluated recognition memory through (C) 1 h and (D) 24 h discrimination indices. Data are expressed as mean ± SEM. Statistical analysis was performed using one-way ANOVA followed by Tukey’s post-hoc test. (n = 8 in each group). Significant differences between groups are denoted as **P < 0.01, ***P < 0.001, while non-significant differences are indicated as ns.

### Regulation of proteins involved in Aβ accumulation and clearance through FUS-TCS

Given that Aβ accumulation is a key feature of AD and that FUS-TCS improved behavioral outcomes in mice injected with Aβ, we aimed to examine whether this treatment affects hippocampal Aβ concentrations and the expression of proteins involved in Aβ clearance. As shown in Figure 5A, Aβ-injected mice showed a significant increase in hippocampal Aβ expression, reaching 1.74 times that of the control group. Importantly, FUS-TCS treatment significantly lowered Aβ levels compared to the Aβ-only group, bringing them back to the control level. After measuring Aβ levels, we examined the expression of proteins related to Aβ clearance. Figure 5B shows that hippocampal HSP70 expression was slightly higher in Aβ-injected mice but not significantly different from the control group, whereas FUS-TCS treatment significantly raised HSP70 levels compared to both control and Aβ groups, reaching 1.57 times the control level. We also investigated IDE, a well-known enzyme that degrades Aβ. Previous studies suggest that reduced IDE expression contributes to Aβ accumulation in AD, and that restoration of IDE levels may mitigate this pathology [42,43]. In our study, as shown in Figure 5C, the Aβ group exhibited a significant downregulation of IDE expression, decreasing to 0.63-fold relative to the control group. Notably, FUS-TCS treatment resulted in a substantial recovery of IDE expression, reaching 0.99-fold of the control level with no significant difference. Additionally, NEP, another critical Aβ-degrading enzyme involved in Aβ degradation and clearance [44,45], was also examined. Reduced NEP expression has been linked to Aβ accumulation, while increasing NEP levels may lower Aβ in AD models [46–48]. Our results showed that NEP expression was markedly decreased after Aβ injection, declining to 0.58-fold of the control level, but this reduction was significantly improved by FUS-TCS treatment, as shown in Figure 5D. Collectively, these results indicate that FUS-TCS may influence critical molecular pathways involved in Aβ clearance, which could contribute to the cognitive improvements observed in treated mice.

**Figure 5.**
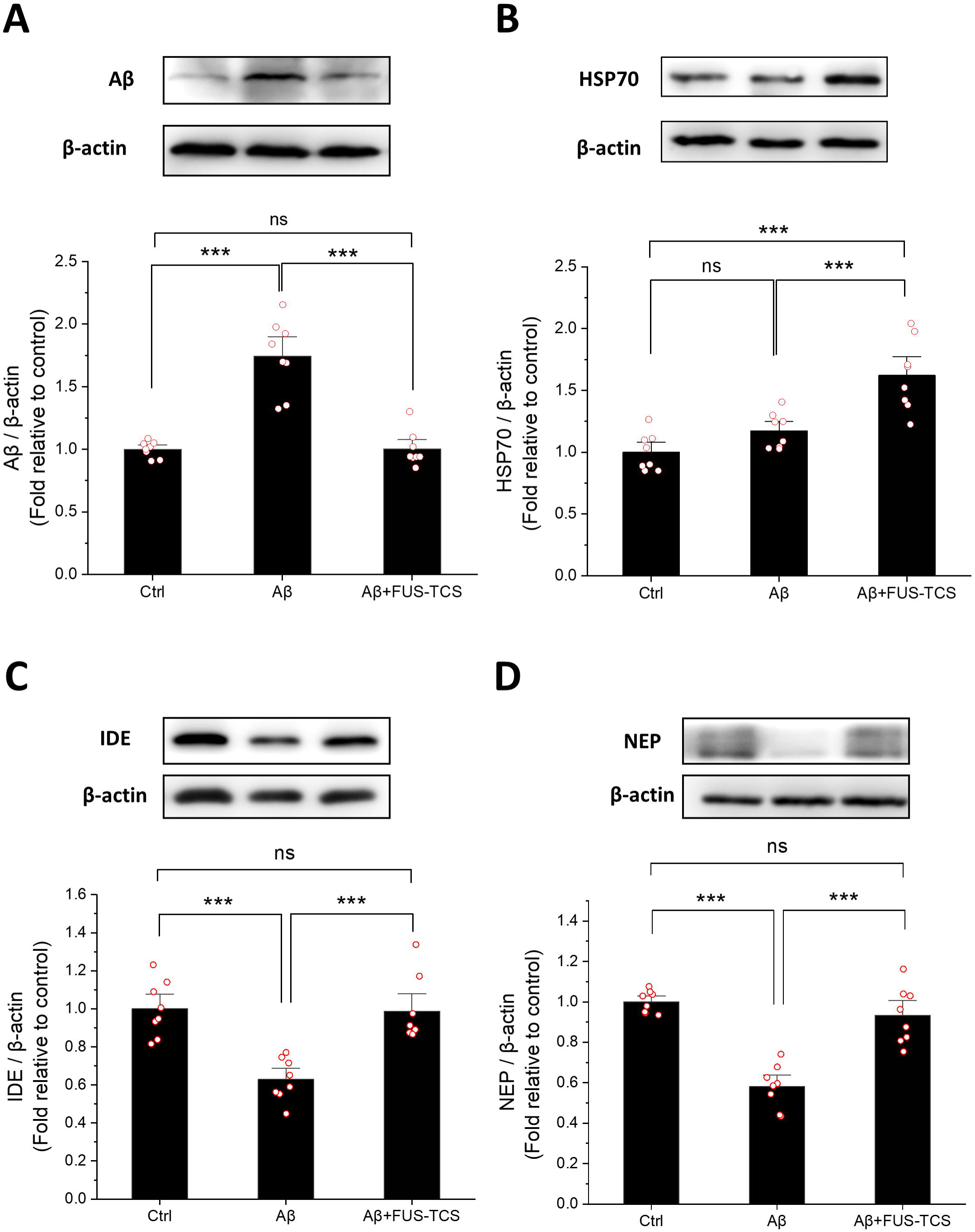
Effect of FUS-TCS treatment on the expression of Aβ, HSP70, IDE, and NEP in the mouse hippocampus. Representative Western blot images and quantification of (A) Aβ, (B) HSP70, (C) IDE, and (D) NEP protein levels in the mouse hippocampus to evaluate the anti-Aβ effect. β-actin was used as an internal control. Relative expression levels were normalized to the control group and presented as fold changes. Data are expressed as mean ± SEM. Statistical analysis was performed using one-way ANOVA followed by Tukey’s post-hoc test. (n = 8 in each group). Significant differences between groups are denoted as **P < 0.01, ***P < 0.001, while non-significant differences are indicated as ns.

### Expression of antioxidant proteins SIRT1 and SOD2 after FUS-TCS treatment

Oxidative stress, driven by elevated reactive oxygen species (ROS), plays a critical role in neuronal injury and the progression of AD, underscoring the therapeutic importance of modulating oxidative stress [49–51]. SIRT1, a NAD⁺-dependent deacetylase, is crucial for regulating cellular defense mechanisms by influencing several downstream pathways involved in antioxidant and anti-inflammatory responses [52,53]. Previous research has shown that SIRT1 levels are decreased in AD patients and inversely correlated with Aβ accumulation [54,55]. Studies also suggest that raising SIRT1 expression is linked to decreased Aβ accumulation and may alleviate AD pathology in animal models [56–58]. In this study, as shown in Figure 6A, SIRT1 expression in the Aβ group was reduced to 0.73-fold relative to the control. This decrease was ameliorated in the group treated with FUS-TCS, where SIRT1 levels did not significantly differ from the control. Additionally, the antioxidant protein SOD2, a well-characterized downstream target of SIRT1-regulated antioxidant pathways, has been shown to increase in response to elevated SIRT1 expression [59–61]. Research also indicates that SOD2 can decrease hippocampal superoxide concentrations and amyloid plaque burden in AD mice [62]. Consistent with these findings, Figure 6B demonstrates that SOD2 expression was significantly decreased to 0.60-fold of the control level in the Aβ group, whereas FUS-TCS treatment restored SOD2 expression to levels comparable to the control, with no significant difference observed. Collectively, these results indicate that FUS-TCS treatment reinstates the expression of antioxidant proteins SIRT1 and SOD2 in the hippocampus, suggesting a potential role in reducing oxidative stress.

**Figure 6.**
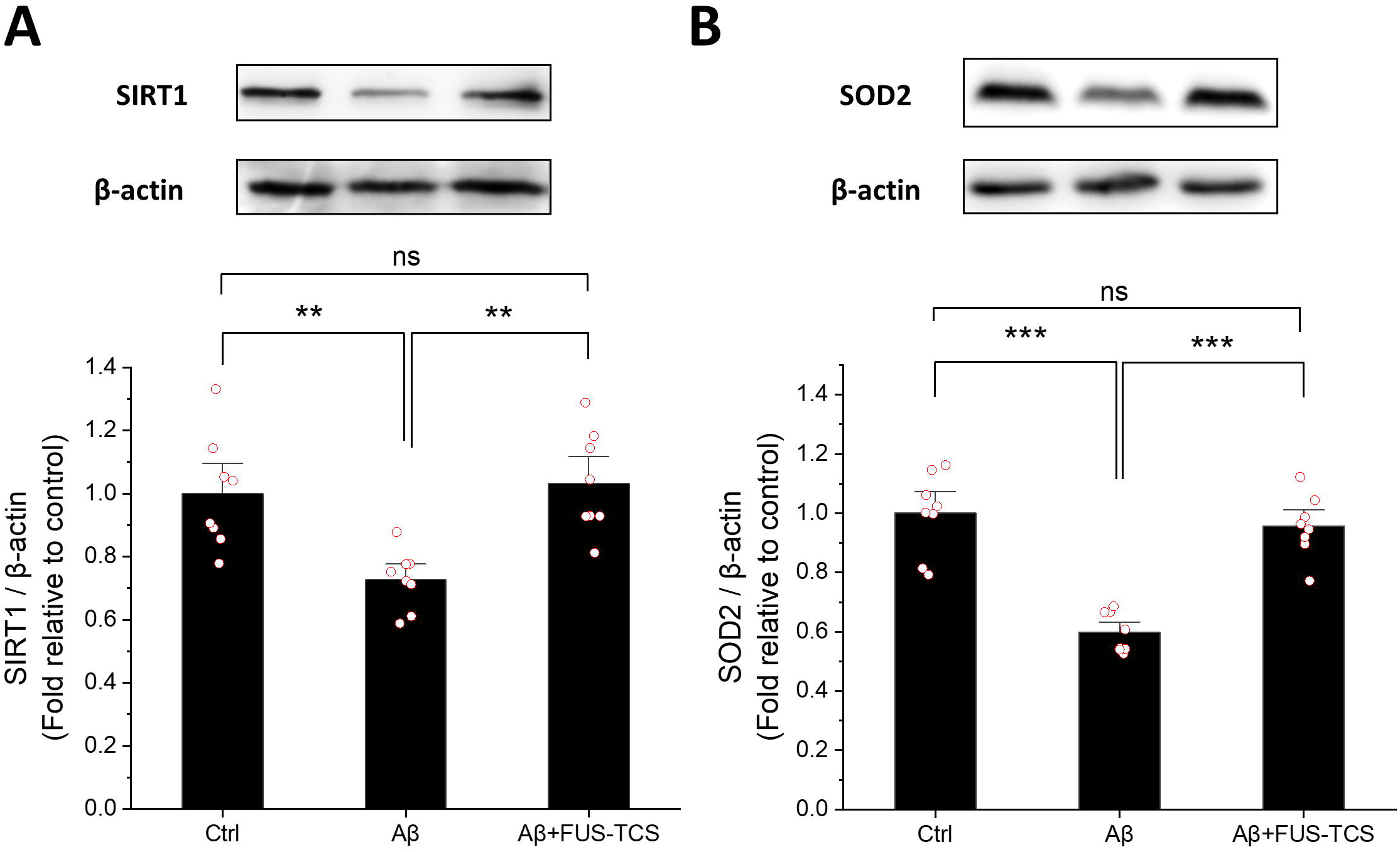
Effects of FUS-TCS treatment on the expression of antioxidant proteins SIRT1 and SOD2 in the mouse hippocampus. Representative Western blot images and quantitative analyses of (A) SIRT1 and (B) SOD2 protein levels in the hippocampus to evaluate antioxidant effects. β-actin was used as an internal control. Relative expression levels were normalized to the control group and presented as fold changes. Data are expressed as mean ± SEM. Statistical analysis was performed using one-way ANOVA followed by Tukey’s post-hoc test. (n = 8 in each group). Significant differences between groups are denoted as **P < 0.01, ***P < 0.001, while non-significant differences are indicated as ns.

## Discussion

AD is a leading cause of dementia, and as the population ages, the threat it poses to humanity is becoming increasingly serious. Researchers are focused on finding effective treatments, with the primary approach being the removal of Aβ [63,64]. However, the BBB makes drug delivery extremely challenging, and the side effects of treatments also present significant obstacles [65,66]. Consequently, the exploration of alternative therapeutic modalities is of essential importance.

Hyperthermia has emerged as a promising non-pharmacological intervention, given that controlled thermal stimulation can promote neuroprotection [67,68]. Nonetheless, the challenge remains in delivering thermal energy specifically to cerebral tissues and activating corresponding neuroprotective signaling pathways. Unlike whole-body hyperthermia, our prior research employed localized thermal stimulation to the cranial region of murine models using polyimide thermofoil, resulting in amelioration of recognition deficits [37]. In the current study, we advance this approach by administering thermal stimulation directly into the brains of mice using FUS, with the objective of modulating protein expression and mitigating cognitive impairments.

To prevent damage from excessive heat exposure, our team has developed an optimized thermal stimulation method, which we refer to as TC-HT. This technique preserves the therapeutic advantages of thermal stimulation through intermittent heat cycling while minimizing cumulative thermal stress. Our previous studies have confirmed the safety of TC-HT and demonstrated that it is more effective than traditional HT, both in vitro and in vivo [35–37]. This study incorporates the TC-HT concept through the application of FUS technique, which is termed as FUS-TCS. Importantly, the use of FUS-TCS did not adversely impact the general health or exploratory behaviors of mice, as assessed by the Y-maze and NOR tests. Remarkably, FUS-TCS enhanced spontaneous alternation performance in the Y-maze, whereas mice subjected to continuous FUS-HTS exhibited comparatively diminished success rates (Figure 2). These findings substantiate the behavioral advantages of FUS-TCS and affirm its safety as an in vivo neuromodulatory approach.

Given that cognitive impairment is a defining characteristic of AD, we subsequently assessed the impact of FUS-TCS on behavioral performance in mice with Aβ-induced cognitive deficits. The findings showed that FUS-TCS treatment significantly improved the spontaneous alternation index in the Y-maze test relative to the Aβ-treated group (Figure 3). Moreover, in the NOR test, FUS-TCS effectively mitigated Aβ-induced cognitive impairments, as evidenced by enhancements in both short-term and long-term memory assessments (Figure 4). These in vivo results suggest that FUS-TCS holds promise as a therapeutic approach for ameliorating memory deficits and cognitive dysfunction associated with Aβ pathology.

The clearance of Aβ has consistently represented a primary target in AD therapeutic research. This study demonstrated that FUS-TCS can ameliorate Aβ-induced cognitive deficits in murine model (Figures 3 and 4) and also reduce Aβ expression levels in the hippocampus (Figure 5A). The mechanistic basis for these effects may involve the modulation of protective proteins through thermal stimulation. Among these proteins, HSP70, a major heat shock protein, plays a vital role in proteostasis by facilitating the refolding and degradation of misfolded proteins, and has been reported to reduce Aβ aggregation and enhance its clearance [6,15,69,70]. The study showed that the FUS-TCS-treated group exhibited a significant upregulation of HSP70 expression in the hippocampus relative to the Aβ-only group (Figure 5B). In addition to HSP70, Aβ-degrading enzymes such as IDE and NEP are also important for clearing Aβ [12,17]. IDE, in particular, has attracted considerable attention in AD research, with both in vitro and in vivo evidence supporting its role in mitigating Aβ accumulation [46,71–73]. Interestingly, IDE shares functional characteristics with HSPs and its expression increases after heat stimulation [22]. Our previous work also demonstrated that thermal stimulation via polyimide thermofoil applied to the murine head enhances IDE expression in the hippocampus [37]. In this study, we found that FUS-TCS application significantly restored the Aβ-induced reduction in IDE levels (Figure 5C), further underscoring its therapeutic potential in AD. Similarly, NEP, another pivotal Aβ-degrading enzyme expressed in the brain, has been implicated in AD pathogenesis; its decreased expression correlates with Aβ accumulation, whereas overexpression facilitates Aβ degradation [46–48,71,72]. Here, our findings reveal that thermal stimulation via FUS-TCS effectively restored NEP expression levels that were reduced by Aβ (Figure 5D). Overall, these results suggest that FUS-TCS promotes the upregulation of proteins involved in Aβ clearance, thereby enhancing its capacity to mitigate AD-related neuropathology.

The regulation of oxidative stress represents a critical focus within AD research. SIRT1, a NAD⁺-dependent deacetylase, has been identified as a major regulator of oxidative stress, functioning to reduce ROS accumulation and thereby exert neuroprotective effects [74,75]. Research has shown that SIRT1 levels are decreased in individuals with AD [55,76,77]. On the other hand, upregulation of SIRT1 has been found to attenuate oxidative stress and contribute to the reduction of Aβ accumulation [58,78]. In this study, our findings revealed that application of FUS-TCS resulted in an upregulation of SIRT1 levels (Figure 6A). Moreover, SIRT1 is believed to enhance the expression of SOD2, a critical mitochondrial enzyme responsible for scavenging superoxide radicals, thereby strengthening antioxidant defenses. Concomitant increases in both proteins have been observed in cells with greater resistance to oxidative stress [61,79]. Our findings also showed that FUS-TCS treatment elevated SOD2 expression (Figure 6B), supporting the therapeutic potential of this intervention to mitigate oxidative stress in an AD mouse model. Collectively, these results underscore the capacity of FUS-TCS to enhance antioxidant protein levels and its potential benefits for AD treatment.

Thermal stimulation influences biological systems by modulating temperature, thereby affecting cellular functions and critical signaling pathways. This modality has demonstrated efficacy in pain management and cancer treatment, with accumulating evidence indicating its potential neuroprotective effects [68,80–82]. A pivotal consideration in applying thermal stimulation to the brain is the capacity to deliver localized heating with precise thermal dose control to avoid the risk of adverse effects. In our prior research, thermal energy was administered to murine heads via a polyimide thermofoil, and the findings showed that TC-HT enhanced cognitive performance [37]. However, this technique relies on indirect brain heating through conduction rather than direct targeting [37]. Due to the small size and thin cranial structure of mice, this method effectively affects brain proteins, though optimization remains possible using other approach. To improve upon this, the current study employs FUS as a thermal stimulation modality, utilizing its capacity for focal energy delivery to directly and precisely regulate local temperature in the brain. To prevent excessive heat accumulation, we incorporated the TC-HT approach, which includes intermittent cooling intervals during thermal stimulation. This cyclic approach lowers the risk of thermal damage associated with traditional HT by introducing rest and cooling intervals, thereby preserving the therapeutic benefits of thermal stimulation while enhancing safety (Figure 2). Our results showed that FUS-TCS application improved the spatial cognition and memory impairments in vivo (Figures 3 and 4). At the molecular level, FUS-TCS significantly upregulated several proteins implicated in Aβ clearance, including the molecular chaperone HSP70 and the Aβ-degrading enzymes IDE and NEP (Figure 5B–D), which may underlie the observed reduction in Aβ levels (Figure 5A). Additionally, FUS-TCS enhanced the expression of SIRT1 and SOD2 (Figure 6A and B), suggesting involvement in oxidative stress regulation that may contribute to cognitive improvements (Figures 3 and 4). In summary, this study offers initial evidence that FUS-TCS is a viable method for delivering localized thermal stimulation to the brain. It may also provide an alternative to traditional drug therapies, potentially overcoming challenges related to BBB permeability and adverse drug effects. Further research should focus on assessing long-term safety, elucidating underlying mechanistic pathways, and exploring clinical applicability. Overall, these findings encourage further exploration of thermal neuromodulation as a promising direction for AD treatment.

## Materials and Methods

### Ethics statement

Animal experiments were conducted according to the regulations approved by the Animal Ethical Committee of National Taiwan University (approval protocol number: NTU-112-EL-00158) Mice were monitored daily for signs of distress, with predefined humane endpoints including reduced social interaction, anorexia, a body weight loss exceeding 15%, hunching posture, or inability to evade handling. Hydrodynamic injections and thermal treatments were administered under anesthesia using 3% isoflurane (Panion & BF Biotech Inc., Taipei, Taiwan), ensuring maximum efforts to reduce any potential discomfort or distress to the animals. For sacrifice, mice were humanely euthanized by carbon dioxide (CO_2_) gas asphyxiation followed by cervical dislocation and were classified as non-survivors.

### Animals

C57BL/6 male mice (6–8 weeks old, 22–26 g) were obtained from the National Laboratory Animal Center. The mice were kept in controlled environmental conditions with regulated temperature and humidity, maintained on a 12 h light/dark cycle (lights on from 7 a.m. to 7 p.m.), and had free access to a standard diet and water. Animal experiments were conducted according to the ethical standards approved by the Animal Ethical Committee of National Taiwan University. The mice were randomly divided into the control, Aβ, FUS-TCS, and FUS-HTS groups. Prior to the application of FUS-TCS or FUS-HTS, the fur on the mice’s heads was shaved, and ultrasound transmission gel (TING YU MEDICAL ENTERPRISE CO., LTD., Taichung, Taiwan) was applied carefully at the interface between the transducer and the animal’s skin to ensure optimal ultrasound transmission.

### Aβ administration

Aβ_25-35_ peptides were aggregated into fibrils by incubating them in a 0.9% sterile saline solution at a concentration of 1 mg/mL, maintained at 37 °C for 7 days. On day 0, 10 μL of this Aβ solution was i.c.v.-injected into the brains of mice in the Aβ, FUS-TCS, and FUS-HTS groups. Meanwhile, mice in the control group received an equivalent volume (10 μL) of sterile saline solution via the same i.c.v. injection procedure.

### Phantom preparation

A bilayer phantom was fabricated using an acrylamide-based hydrogel to preliminarily evaluate the heating performance of FUS. Briefly, the precursor solution was prepared by combining 33% (v/v) of a 30% acrylamide stock solution (CAS No. 79-06-1; BioShop Canada Inc., Burlington, Ontario, Canada) with 61.8% (v/v) deionized water under continuous stirring. Subsequently, 4.5% (v/v) anhydrous glycerol (CAS No. 56-81-5; BioShop, Canada) was incorporated and thoroughly homogenized. Polymerization was initiated by the addition of 0.5% (v/v) of 10% ammonium persulfate (CAS No. 7727-54-0; BIO BASIC, Inc., Canada) and 0.2% (v/v) N,N,N’,N’-tetramethylethylenediamine (TEMED, CAS No. 110-18-9; Sigma-Aldrich, Merck KGaA). To create the bilayer lateral configuration, the precursor solution was evenly divided into two aliquots. One aliquot was supplemented with thermochromic powder (Shenzhen Huancai Color-Changing Technology Co., Ltd.) at a final concentration of 1% (w/v) to produce a white, thermoresponsive region that undergoes a color change at 40 °C. A temporary divider was positioned within the mold to spatially separate the two compartments, and each formulation was cast simultaneously into its respective section. Following partial gelation, the divider was carefully removed to permit interfacial fusion. The phantom was then left to fully polymerize under ambient conditions, resulting in a bilayer structure comprising adjacent transparent and thermochromic domains.

### Applications of FUS-HTS and FUS-TCS

Ultrasound sonication was produced using a single-element focused piezoelectric transducer operating at 1.2 MHz, with a diameter and radius of curvature both measuring 20 mm. A power meter/sensor module (Bird 4421, Bird Electronic Corp., Cleveland, OH) was used to measure the input electrical power. The ultrasound signal was generated by a function generator (SG382; Stanford Research Systems, Sunnyvale, CA, USA) and amplified by a power amplifier (25A250, Amplifier Research, Souderton, PA, USA). To monitor the actual temperature at the focal point, a K-type thermocouple was inserted into the brain tissue of anesthetized mice during both FUS-TCS and FUS-HTS treatments. For the FUS-TCS procedure, the right cerebral hemispheres of the mice underwent a 10-cycle repeated treatment, where each cycle consisted of 3 min FUS heating activation followed by a 1 min rest interval. On the other hand, the FUS-HTS group received continuous FUS heating treatment on the right brain hemisphere for 30 min without interspersed rest intervals. The temperature increase induced by FUS heating in a phantom caused visible color changes, as shown in Figure 1A and B, while the monitored temperatures of the mice’s brains during FUS-TCS and FUS-HTS applications were presented in Figure 2A.

### Experimental schedule design

Mice were randomly assigned to various treatment groups, including control, FUS-TCS, FUS-HTS, Aβ, Aβ+FUS-TCS, and Aβ+FUS-HTS groups. On day 0, 10 μL of Aβ solution was i.c.v.-injected into the brains of mice in the Aβ, Aβ+FUS-TCS, and Aβ+FUS-HTS groups. Meanwhile, control, FUS-TCS, and FUS-HTS groups were administered an equivalent volume (10 μL) of phosphate buffered saline (PBS, HyClone; GE Healthcare Life Sciences, Chicago, IL, USA) via i.c.v. injection to serve as controls. Thermal treatments were administered on days 4, 8, and 12 to mice in the FUS-TCS, FUS-HTS, Aβ+FUS-TCS, and Aβ+FUS-HTS groups, as shown in Figure 2A. Subsequently, locomotor activity, memory, and recognition were evaluated using Y-maze and NOR tests, which rely on the natural curiosity of mice to explore new environments or objects [38–41]. The Y-maze test was conducted on day 13, while the NOR tests, assessing short-term (1 h) and long-term (24 h) memory, were performed on days 14 and 15, respectively.

### Y-maze test

The Y-maze test was utilized to assess the spatial working memory of mice. The maze comprised three identical arms (40 cm × 13 cm × 6.5 cm) interconnected at a central junction, with each arm separated by an angle of 120°. The mouse under assessment, which had no prior exposure to the maze, was placed at the end of the starting arm and allowed to freely explore all three arms for a duration of 8 min to measure spontaneous alternation. This test relies on the natural curiosity of mice to sequentially explore all three arms in a maze without external stimuli, making it a reliable indicator of spatial working memory [38,39]. After each assessment, olfactory cues in the maze were eliminated by cleaning the maze with 70% ethanol. The mouse’s behavior was recorded, noting the order of entries into the three arms labeled ‘’A’’, ‘’B’’, and ‘’C’’. An arm entry was defined as the mouse’s hind paws fully entering an arm. A correct alternation was identified when the mouse consecutively entered all three arms in overlapping triplets, such as A-B-C, A-C-B, B-A-C, B-C-A, C-A-B, and C-B-A. (for example, the sequence of A-C-B-A-C contains three correct alternations) [38,39]. The spontaneous alternation index was calculated as (number of correct alternations) / (total number of arm entries – 2) × 100. The total number of arm entries was also recorded as a measure of the mouse’s locomotor activity.

### NOR test

The NOR test was employed to evaluate the recognition memory of the animals. The NOR test consists of a habituation phase, a familiarization phase and a test phase. During habituation, mice were placed individually in an empty box (40 cm × 40 cm × 30.5 cm) for 8 min. In the familiarization phase, the assessed mouse was placed in the same box for 8 min with two identical objects ‘’X’’ positioned near two diagonal corners. Short-term memory was assessed 1 h after familiarization by returning the mouse to the box for the test phase, where one of the original objects was replaced with a novel object ‘’Y’’ of a different shape. The mouse’s behavior was recorded with an overhead video camera for 8 min. Long-term memory was evaluated 24 h later in a similar manner, but object ‘’Y’’ was replaced with another novel object ‘’Z’’. After each phase, the box was cleaned with 70% ethanol to remove olfactory cues. Exploration was defined as the mouse sniffing or manipulating an object with its nose or forepaws. The discrimination index was calculated as (Tn-Tf) / (Tn+Tf), where Tn and Tf represent the time spent exploring the novel and familiar objects, respectively.

### Brain tissue collection and Western blot sample preparation

Following the NOR tests on day 15, mice were euthanized using CO_2_ inhalation. After confirming the death, their brains were quickly extracted and rinsed with PBS supplemented with 1% fresh protease inhibitor cocktail (EMD Millipore, Billerica, MA, USA). The entire hippocampi were rapidly dissected on an ice-cold plate, and kept in ice-cold RIPA lysis buffer (MilliporeSigma, Burlington, MA, USA) supplemented with 1% fresh protease inhibitor cocktail. The hippocampal tissues were then homogenized using ultrasound treatment (three cycles of 5 s rectangular pulses with 5 s intervals at 50% amplitude, using a Q125 sonicator; Qsonica, Newton, CT, USA) and centrifuged at 23000×g for 30 min at 4 °C. After centrifugation, the supernatants were collected, and protein concentrations were quantified using the Bradford protein assay (BioShop Canada Inc.). Protein samples were stored at −80 °C for subsequent Western blot analyses.

### Western blot analysis

Proteins extracted from homogenized hippocampi (20 µg) were separated using sodium dodecyl sulfate polyacrylamide gel electrophoresis (SDS-PAGE) and then transferred onto polyvinylidene difluoride (PVDF) membranes (MilliporeSigma). The membranes were blocked for 1 h at room temperature using 5% bovine serum albumin (BSA) in Tris-buffered saline containing 0.1% Tween-20 (TBST) (BioShop Canada Inc.). Following blocking, the membranes were incubated overnight at 4 °C with primary antibodies diluted in blocking buffer, then followed by three rinses with TBST. The primary antibodies used in this study included anti-Aβ (cat. no. sc-28365; Santa Cruz Biotechnology, Inc., Dallas, TX, USA), anti-IDE (cat. no. ab32216; Abcam, Cambridge, UK), HSP70 (cat. no. 4872; Cell Signaling Technology, Inc.), anti-NEP (cat. no. GTX111680; Gentex, Irvine, CA, USA), anti-SIRT1 (cat. no. 9475; Cell Signaling Technology, Inc.), anti-SOD2 (cat. no. 13141; Cell Signaling Technology, Inc.), and anti-β-actin (cat. no. GTX110564; Gentex). After washing with TBST, membranes were incubated with horseradish peroxidase-conjugated goat anti-mouse secondary antibody (for detecting anti-Aβ primary antibody; cat. no. ab205719; Abcam) or goat anti-rabbit secondary antibody (for the other primary antibodies; cat. no. AB_2313567; Jackson ImmunoResearch Laboratories, Inc., West Grove, PA, USA) for 1 h at room temperature. β-actin was used as the loading control to normalize protein levels. All the antibodies were diluted according to the manufacturers’ recommended optimal concentrations. Protein bands were detected using an enhanced chemiluminescence substrate (Advansta, San Jose, CA, USA) and imaged with an Amersham Imager 600 system (AI600; GE Healthcare Life Sciences). Quantitative analysis of protein band intensities was performed using Image Lab software (version 6.0.1, Bio-Rad Laboratories, Inc.).

### Statistical analysis

All statistical data were presented as mean ± SEM and analyzed using one-way analysis of variance (ANOVA) followed by Tukey’s post hoc test in OriginPro 2022 software (OriginLab Corporation, Northampton, MA, USA). *P*-value < 0.05 indicated a statistically significant difference.

## Conclusions

This study reveals that FUS-TCS, which administers mild, localized, and intermittent hyperthermia to the brain, significantly mitigates Aβ-induced cognitive impairments in mice. The treatment induces upregulation of HSP70 and enhances the activity of Aβ-degrading enzymes, including IDE and NEP. Additionally, it elevates the expression of oxidative stress regulators such as SIRT1 and SOD2, indicating a multifaceted neuroprotective mechanism. These results support FUS-TCS as a safe and promising non-pharmacological approach for AD treatment. Future research is warranted to assess the long-term safety, durability of cognitive improvements, and the underlying biological mechanisms across various AD models to determine its translational applicability.

## Acknowledgements

We thank the Animal Resource Center and the Consortium of Integrative Biomedical Science Key Technology at National Taiwan University for their technical support.

## Data Availability Statement

The data presented in this study are available upon request from the corresponding author.

## Author Contributions

Conceptualization, C.-Y.C.; methodology, G.-B.L., H.-H.L. and C.-Y.C.; validation, G.-B.L., H.-H.L., Y.-Y.K., Y.-M.C., F.-T.H., Y.-W.W., Y.K., C.C. and C.-Y.C.; formal analysis, G.-B.L., H. H.L., Y.-Y.K., Y.-M.C. and C.-Y.C.; investigation, G.-B.L., H.-H.L. and C.-Y.C.; resources, C.-Y.C.; data curation, G.-B.L., H.-H.L., Y.-Y.K., Y.-M.C., F.-T.H., Y.-W.W., Y.K., C.C. and C.-Y.C.; writing—original draft preparation, G.-B.L., and C.-Y.C.; writing—review and editing, G.-B.L. and C.-Y.C.; supervision, C.-Y.C.; funding acquisition, C.-Y.C. All authors have read and agreed to the published version of the manuscript.

## Institutional Review Board Statement

All experimental procedures adhered to the established guidelines of the Animal Ethical Committee of National Taiwan University (approval protocol number: NTU-112-EL-00158 (11 November 2024)).

## Informed Consent Statement

Not applicable.

## Abbreviations

AD: Alzheimer’s disease
Aβ: β-amyloid
BBB: blood-brain barrier
HT: hyperthermia
FUS: focused ultrasound
HIFU: high-intensity focused ultrasound
TC-HT: thermal cycling-hyperthermia
FUS-TCS: focused ultrasound-mediated thermal cycling stimulation
FUS-HTS: focused ultrasound-mediated hyperthermia stimulation
i.c.v.: intracerebroventricular
NOR: novel object recognition
HSP70: heat shock protein 70
IDE: insulin-degrading enzyme
NEP: neprilysin
SIRT1: sirtuin 1
SOD2: superoxide dismutase 2
ROS: reactive oxygen species
ANOVA: analysis of variance

